# Atrous Convolution with Transfer Learning for Skin Lesions Classification

**DOI:** 10.1101/746388

**Authors:** Md. Aminur Rab Ratul, Mohammad Hamed Mozaffari, Enea Parimbelli, WonSook Lee

## Abstract

Skin cancer is a crucial public health issue and by far the most usual kind of cancer specifically in the region of North America. It is estimated that in 2019, only because of melanoma nearly 7,230 people will die, and 192,310 cases of malignant melanoma will be diagnosed. Nonetheless, nearly all types of skin lesions can be treatable if they can be diagnosed at an earlier stage. The accurate prediction of skin lesions is a critically challenging task even for vastly experienced clinicians and dermatologist due to a little distinction between surrounding skin and lesions, visual resemblance between melanoma and other skin lesions, fuddled lesion border, etc. A well-grounded automated computer-aided skin lesions detection system can help clinicians immensely to prognosis malignant skin lesion in the earliest possible time. From the past few years, the emergence of machine learning and deep learning in the medical imaging has produced several image-based classification systems in the medical field and these systems perform better than traditional image processing classification methods. In this paper, we proposed a popular deep learning technique namely atrous or, dilated convolution for skin lesions classification, which is known to have enhanced accuracy with the same amount of computational cost compared to traditional CNN. To implement atrous convolution we choose the transfer learning technique with several popular deep learning architectures such as VGG16, VGG19, MobileNet, and InceptionV3. To train, validate, and test our proposed models we utilize HAM10000 dataset which contains total 10015 dermoscopic images of seven different skin lesions (melanoma, melanocytic nevi, Basal cell carcinoma, Benign keratosis-like lesions, Dermatofibroma, Vascular lesions, and Actinic keratoses). Four of our proposed dilated convolutional frameworks show promising outcome on overall accuracy and per-class accuracy. For example, overall test accuracy achieved 87.42%, 85.02%, 88.22%, and 89.81% on dilated VGG16, dilated VGG19, dilated MobileNet, and dilated IncaptionV3 respectively. These dilated convolutional models outperformed existing networks in both overall accuracy and individual class accuracy. Among all the architectures dilated InceptionV3 shows superior classification accuracy and dilated MobileNet is also achieving almost impressive classification accuracy like dilated InceptionV3 with the lightest computational complexities than all other proposed model. Compared to previous works done on skin lesions classification we have experimented one of the most complicated open-source datasets with class imbalances and achieved better accuracy (dilated inceptionv3) than any known methods to the best of our knowledge.

## 1 Introduction

Day by day the occurrence of skin cancer has spread shockingly, and it is one of the most common forms of cancer in the world [1, 2]. It is the most pervasive type of cancer in the USA with dreadfully 5 million cases happening per year [3, 4, 5]. There are various kinds of skin lesions such as intraepithelial carcinoma, melanoma, squamous cell carcinoma, basal cell carcinoma, etc. [6, 7, 8]. Among all of them, melanoma is extremely cancerous and causes the extensive amount of deaths annually (over 9000 deaths in the USA in 2017) [9]. Early diagnosis can cure near 95% of melanoma, however, if the lesion reaches in lymphatics then the survival rate falls down to nearly 62% [10]. The treatment accuracy of dermoscopy can vary between 75%-84% [11, 13, 14].

The unaided manual detection system of skin lesions detection can achieve the accuracy of about 60%, however, it is human-labor intensive [11]. It is a noninvasive procedure of skin imaging which needs illuminated and magnified images of skin to increase the spots clarity [12] and effacing the surface reflection improves further the visual effect of skin spots. To observe malignant melanoma in the early stage, ABCD-rule (Asymmetry, Border, Irregularity, Color variation and Diameter) is the most popular technique among them and other procedural algorithms such as Menzies method, 3-point checklist, and 7-point checklist have been developed to boost the scalability of dermoscopy [13, 15]. However, many clinicians do not rely on these methods and they heavily rely on their personal experience [16]. The dermoscopy imaging technique needs vast visual exploration, similarities, and dissimilarities between separate lesions, and the necessity of follow-ups over years create a difficult diagnosis situation for the experts and these procedures are more prone to mistake.

Due to these reasons and to detect any skin lesion in the earliest possible stage researchers has proposed many computerized lesion detection models. Recently, deep learning model has made magnificent advancement in computer vision, started with AlexNet [20] in 2012. Then, a large number of deep learning architecture have been developed to detect, classified, and to diagnosis different types of diseases through medical imaging [21]. Mostly they are supervised machine learning methods requiring annotated training data and last few years dermoscopy has been able to produce an unprecedently huge amount of well-annotated images of skin lesions which help these automated systems vigorously [17, 18, 19]. The most popular deep convolutional neural networks (CNN) not only use to predict the specific kind of diseases but also utilize to differentiate the classes of illness [22]. Therefore, automated deep learning based medical imaging system can be a useful tool to help the dermatologist to focuses on different areas such as skin lesions detection, segmentation, and classification.

In this paper, we propose to apply a new approach namely atrous (or dilated) convolution to classify multi-class skin lesion. Atrous is known to be good for improving accuracy with the same amount of computational complexities compared to traditional CNN. Firstly, we choose four well-known models of deep learning specifically VGG16, VGG19, MobileNet, InceptionV3. Secondly, to get better performance and accuracy we implement atrous convolutional layers in the place of traditional convolutional layers of these architectures. Additionally, we use fine tuning strategy to train every proposed model. To train, validation, and test the models we use HAM10000 dataset which acquired by ViDIR group of Medical University of Vienna [23]. This dataset contains a total of 10015 dermatoscopic images of seven different categories of skin lesions. These seven different categories are melanoma, melanocytic nevi, Basal cell carcinoma, Benign keratosis-like lesions, Dermatofibroma, Vascular lesions, and Actinic Keratoses. Among these lesions, melanoma is the most dangerous one, Actinic keratoses, and Basal cell carcinoma can be cancerous, the rest skin lesions are benign.

The rest of the paper organized as follows: in section 2, discuss some related work. The dataset and methodology illustrate in section 3. Section 4 comprises the performance analysis of the experiment and discussion. Finally, the conclusion and future work presented in section 5.

## 2 Related Work

In the past, many research teams around the world proposed computer-aided algorithms to resolve the issue of skin cancer detection. Most of the algorithms mainly focused on supervised learning, region splitting and merging, thresholding, clustering, and active contour model. Every model has some advantages and also shortcomings [24, 25, 26].

Some recent effort to tackle this problem is as follows. Celebi et al. [27] developed an ensemble four separate thresholding methods to detect skin lesion border. Sigurdsson et al. [29] presented a highly automated and probabilistic method to classify skin lesion where they suggested a feature extraction model for Raman Spectra and a powerful feedforward neural network as a classifier. Barata et al. [30] discuss another approach using two distinct methods namely local features and global features to detect melanoma from dermoscopy images. In the local feature method, they utilize BoF (Bag of Features) classifier for object recognition while in the global method, they use wavelets and segmentation, linear filters or Laplacian pyramids with gradient histogram to extract features like color, shape, and texture from an entire skin lesion. Their claim is that color features exhibit much superior result than texture features alone. In [31] She et al. utilize features like color variation (blue, red, green), diameter, border, and asymmetry to do the classification of melanoma skin lesions. Meanwhile, several pre-processing criteria, for example, lesion localization, contrast enhancement, artificial removal, color transformation, and space transformation are mentioned to accomplish the skin lesion segmentation process [32].

More recently many deep learning methods play a vital role to detect several kinds of skin cancer. Several numbers of deep neural network architectures have been built for classification, segmentation, and detection. Hekler et al. [63] implemented a deep learning architecture for histopathologic diagnosis of melanoma and compare their classification result with experienced histopathologist. The objective of this work to provide a deep learning model for melanoma diagnosis to augment the human assessment. Esteva et al. [34] were utilizing their own labeled dataset by a dermatologist which containing 129,450 clinical images and, in this dataset, they had 3374 dermoscopy images. Furthermore, the whole dataset contains nine classes of skin diseases. The research team use ImageNet pre-trained inceptionv3 and get 72.1 ± 0.9% (mean ± s.d.) overall classification accuracy and two dermatologists obtain 65.56% and 66.0% accuracy on the validation set.. Harangi et al. [35] use an ensemble of DCNN (deep convolutional neural network) to enhance the accuracy of the individual class of ISBI 2017 [62] dataset (contains three classes melanoma, nevus, and seborrheic keratosis). Additionally, to gain high accuracy, they combine classification layers outcome of four separate DCNN models. Rather than training Convolutional Neural Network (CNN) from scratch Kawahara et al. [36] attempted to employ pre-trained ConvNet as their feature extractor and they classify 10 classes of non-dermoscopic skin images.

Bovik and Xie [28] proposed a supervised learning method for skin lesion segmentation where the genetic algorithm merged with a self-generating convolutional neural network (CNN). In [33] Yuan et al. present a 19-layer deep CNN for skin lesion segmentation which is trained end-to-end, and this architecture does not rely on previous knowledge of datasets. An unsupervised learning architecture namely Independent Histogram Pursuit (IHP) proposed by Gomez et al. [37] for the skin lesion segmentation. Further, this model tested on five separate dermatological datasets, and they attain almost 97% segmentation precision. Hybrid thresholding procedure and optimal color channels implemented by Garnavi et al. [38] for an automated segmentation process of skin lesion. Codella et al. presented a hybrid approach in [39] fusing support vector machine (SVM), sparse coding, and deep learning method for melanoma detection in dermoscopy images. Lately, in a new research Codella and his team construct a new automated method merging machine learning and deep learning architectures for the classification and segmentation of skin lesion [40]. In [41] Yu et al. proposed a very deep residual convolutional neural network to differentiate non-melanoma images from melanoma images.

There are several interesting deep learning models which can produce a very impressive result in classification and recognition for dermoscopy images. However, to build this futuristic model for medical imaging and skin lesion detection there is still a lot of places exist where we can make improvement. As one of the most popular deep learning techniques currently applied, atrous convolution is used in various applications like visual question answering [49], optical flow [50], instance segmentation [51], object detection [52] [53], and audio generation [54]. In this paper, we proposed a classification method for skin lesion classification using namely atrous or, dilated convolution with fine tuning technique.

## 3 Dataset and Methods

### 3.1 Dataset

**Figure 1:**
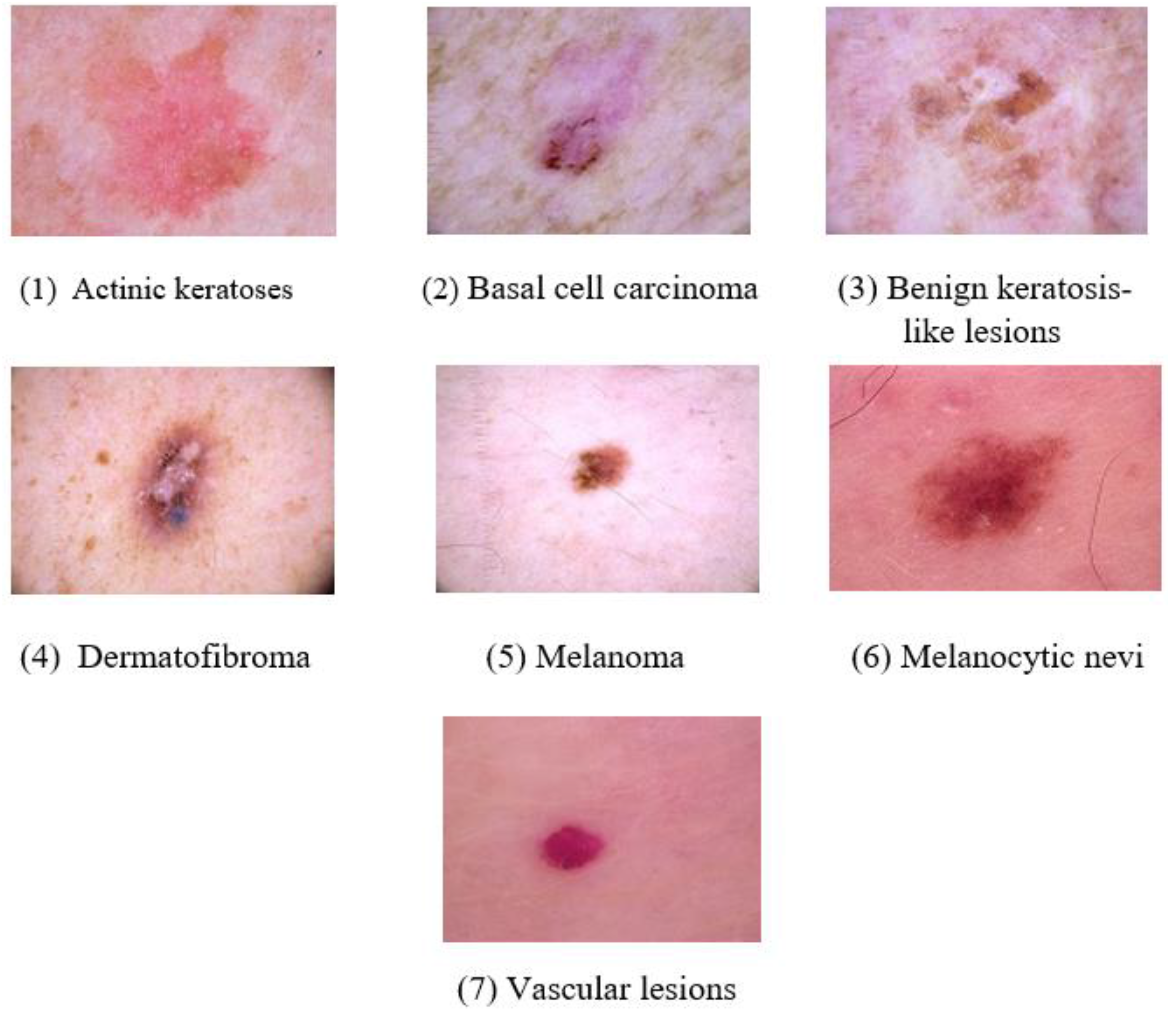
Skin lesions from HAM10000: here Melanoma is extremely dangerous skin lesion, Basal cell carcinoma, and Actinic keratosis can be cancerous. Other skin lesions are can be counted as benign skin lesions

For this research work, we use dermatoscopic skin lesion images from HAM10000 (“Human Against Machine with 10,000 training images”) [23][42]. The dataset consists of a total of 10,015 dermatoscopic images and altogether there are seven different kinds of skin lesions images. These seven skin lesions are Melanoma (1113 images), Melanocytic nevi (6705 images), Basal cell carcinoma (514 images), Benign keratosis-like lesions (1099 images), Dermatofibroma (115 images), Vascular lesions (142 images), and Actinic keratoses (327 images). So, we further divided this dataset as 8011 training images (80%), 1002 validation images (10%), and 1002 test images (10%). HAM10000 is a class imbalanced dataset so that we follow the stratified technique to split this dataset which means when we split the training, validation, and test set, we try to retain the ratio between classes.

#### 3.1.1 Data Pre-Processing

The dimension of images of HAM10000 dataset is 600 × 450. To reduce the computational cost of the proposed deep learning architecture we rescale and resize the images of this dataset. Directly resizing can contort the shape of the lesions and that’s why, at first, we rescale the images by crop the center part of the images and then proportionally lessen the dimension of the image. We take width to height rate as 4: 3, and inspected several combinations of image sizes such as 320 × 320, 512 × 384, 128 × 96, and 64 × 64. Based on overall system accuracy, space complexity, and time complexity 256 × 192 provided foremost output. However, only for atrous MobileNet model we take image size 224 × 224 because MobileNet only accepts four combinations of static square image shapes ((128 × 128), (160 × 160), (192 × 192), (224 × 224)) [57]. We covered this part with more details in performance analysis section and the outcome has been summarized in Table 10. At last, to change the span of intensity values of the pixels of every sample we normalized our data divided each pixel values of an image by 255.0 to rescale the range of 0-255 for each pixel values to 0-1 range.

#### 3.1.2 Data Augmentation

To achieve good performance from a deep learning model we usually need a sheer amount of data [43], so that data augmentation has been used to produce a good amount of data for the model and provide a class-balanced dataset. Our training set contains 8,011 images and we use 200 iterations (or epochs) for the training phase of the proposed DNN. In each iteration, one training example augmented only one time. Additionally, within each epoch transformed images produced from original images for each mini batch. Overall 8,011 transformed training samples are generated in every epoch. So, for 200 iterations from 8,011 images, we able to produce 1,602,200 (200 × 8011) transformed images. Furthermore, we use the same strategy for our validation set containing 1,002 images. In each iteration, we produce 1,002 transformed images to validate our training set.

Moreover, we utilized a series of geometrical transformations on our training set as a part of data augmentation. To transform each image at first, we do horizontal flipping with the probability of 0.50 and then other 0.50 probability accomplished by vertical filliping. Next, we use rotation with range randomly between 0 degrees to 60 degrees. After that, the object which we want to detect may not be placed the only center of the image. Objects can be off-centered in several different ways. Hence, to handle these off-center objects we use randomly horizontal shifting also knows as width shifting and random vertical shifting well known as height shifting. Additionally, the range of both shifting is 0.20. Then to zooming inside the image randomly, we applied zoom with range (−0.2, +0.2). At last, we employ shear mapping or shear transformation to displaces the image horizontally with range of (*x* + 0.2, *y*) or vertically with (*x, y* + 0.2).

**Figure 2:**
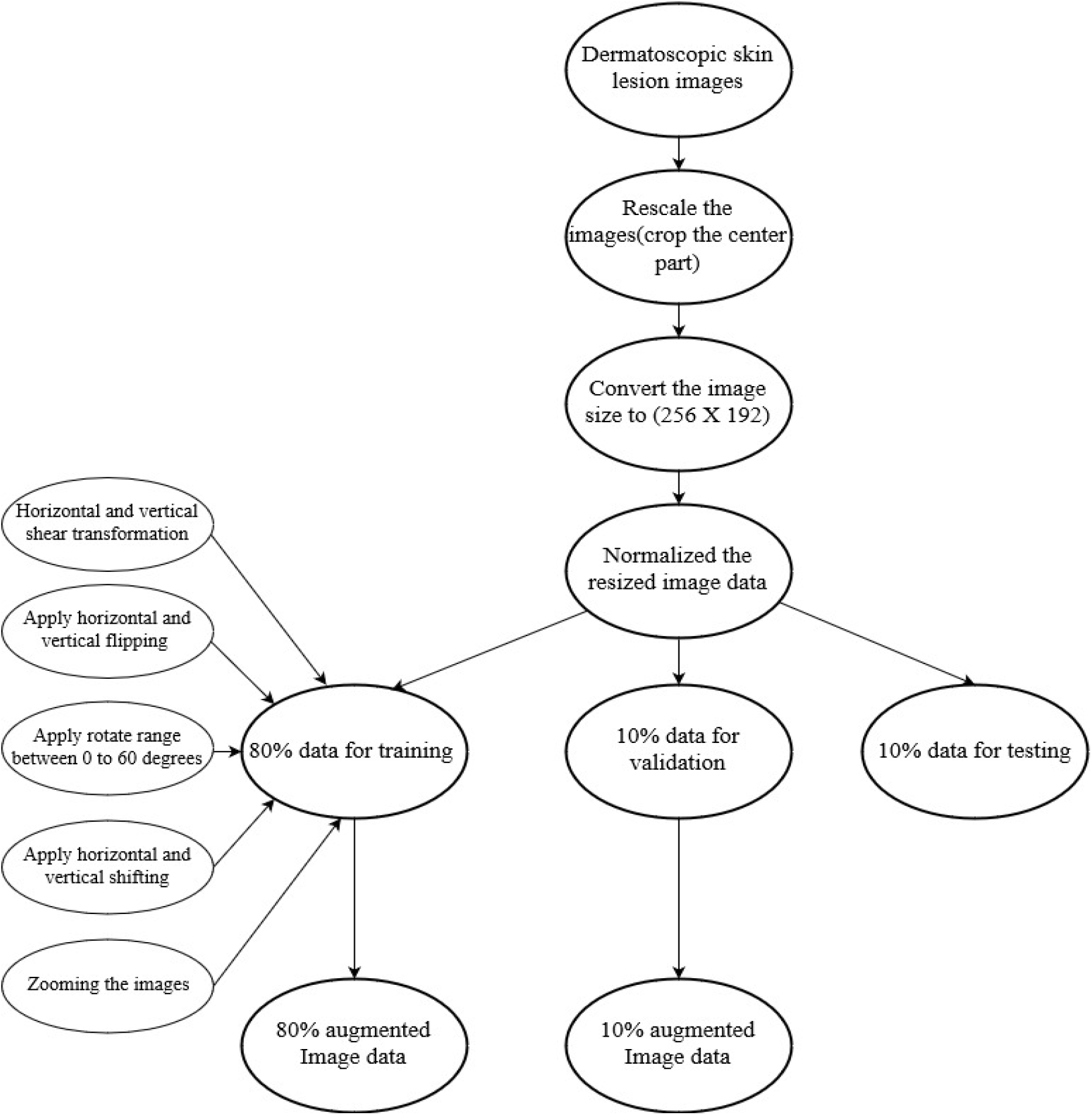
Data pre-processing and data augmentation before feeding into the CNN architectures

### 3.2 Method Description

We aim to help dermatologist to detect any kind of skin lesion as early as possible. In this research work, we choose four state of the art deep learning models such as VGG16 [45], VGG19, MobileNet [46], and InceptionV3 [47] to accomplish our work. On top of the base models, we take a popular approach called atrous or, dilated convolution instead of the conventional convolutional layer. We approach fine tuning technique with dilated convolution for these popular models to bolster the accuracy with the same computational complexity as basic models.

### 3.3 Atrous Convolution

“Atrous” actually comes from a French term called “à trous” which means hole. Originally, researchers invented an algorithm namely “hole algorithm” or “algorithme à trous” for wavelet transformation [48] but right now researchers utilized it for convolution in the deep neural network. “atrous convolutional” is also known as “Dilated Convolution” in the deep learning area.

**Figure 3:**
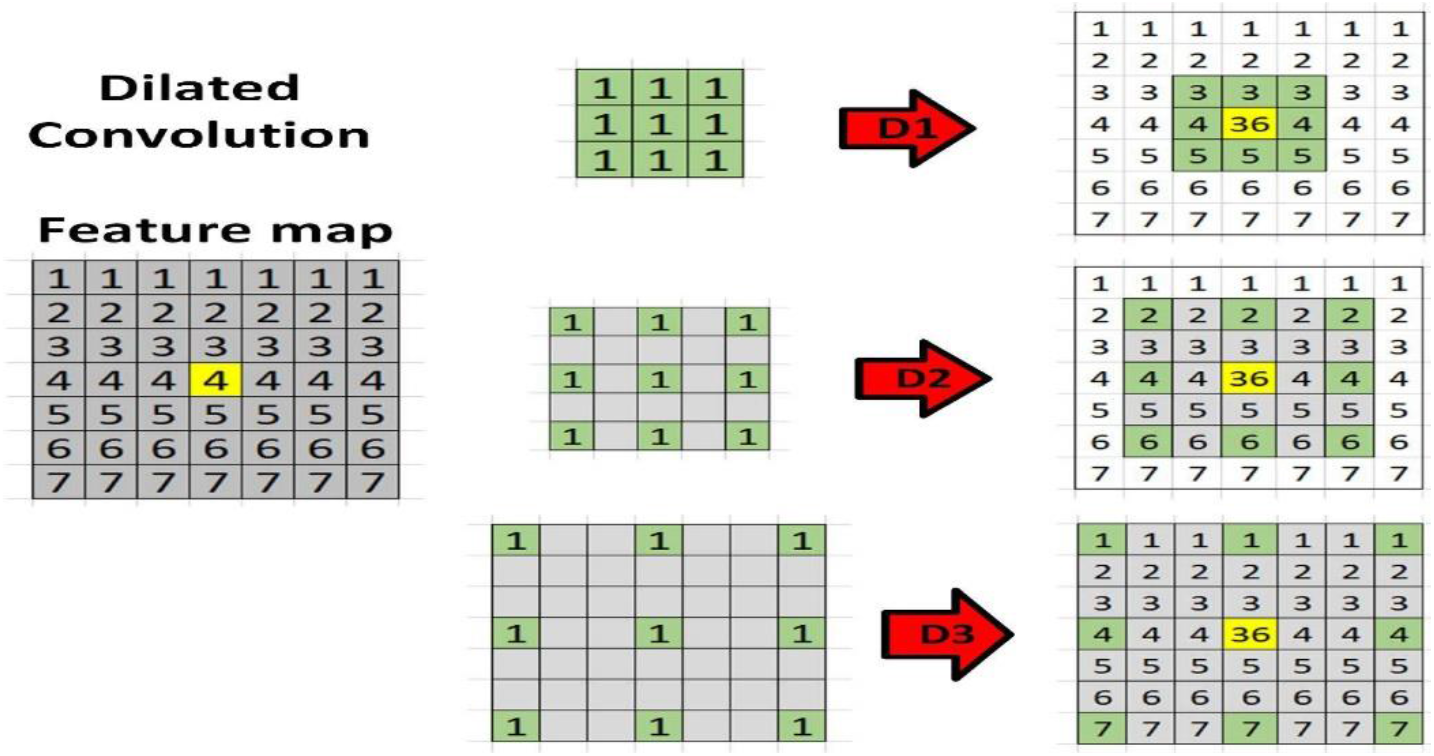
A scenario of dilated convolution for kernel size 3×3. From the top: (a) it is the situation of the standard convolutional layer when the dilation rate is (1,1). (b) when the dilation rate become (2,2) the receptive field increases. (c) in the last case, the dilation rate is (3,3) and the receptive field enlarges even more than situation b

The principle idea of atrous convolution is to insert zeros or “hole” between the kernel of convolutional layers to enhance the image resolution, hence approving dense feature extraction in deep convolutional neural network (DCNNs). Furthermore, atrous convolution used to expand the field of view of the kernel with the identical amount of computational cost. So, it is very much useful with some application which cannot afford larger kernels or, multiple convolutions but needs a broad field of vision.

In one-dimensional scenario, atrous convolution can be defined as:

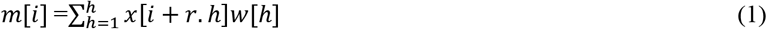

Here, for every location of *i*, *y* is the output and *x[i]* is the input signal (*x* also referred to as a feature map). Moreover, *w[h]* is filter with the length of *h*, and *r* is corresponding to the atrous rate with which we sample *x[i]* (input signal). In the standard convolution r =1 but in the dilated convolution rate of r is always bigger than 1. An instinctive and easy way to comprehend atorus convolution is that inserted r-1 zeros between every two consecutive filters in the standard convolution. If a standard convolution with kernel or filter size *n* × *n*, then the resulting atrous filter or kernel size is *nd* × *nd* where *nd* = *n* + (*n-1*). (*r-1*).

One of the main reasons behind implement this method is without missing any coverage or resolution, atrous convolutions supports exponentially enlarging receptive fields [55]. Let take 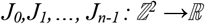 a discrete function and take 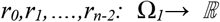 as discrete 3 × 3 kernels or filters. Now, assume employing the filters with exponentially enhancing the atrous rate:

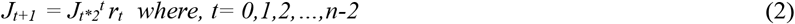

Let, ***k*** the receptive field of an element in ***J_t+1_*** as the sets of elements in ***J_0_*** that adapt the value of ***J_t+1_ (k)***. Let, the extent of the receptive field of ***k*** in ***J_t+1_*** be the number of these components. Therefore, it is suitable to measure the extent of the receptive field of every single component in ***J_t+1_*** will be ***(2^t+2^ −1) × (2^t+2^ −1)***.

#### 3.3.1 Atrous VGG16 and Atrous VGG19 Architecture

VGG16 and VGG19 both have exactly five blocks of standard convolutional layers. To incorporate the larger context view of filters with the exact same number of parameters we change the dilation rate of these standard convolution layers for both the models. When we execute standard convolution and pooling in any model output stride are increasing which means when we go deeper in any model, the output feature map shrinking. It is harmful for any segmentation or classification because in the deep layers spatial or, location information will be lost. On the other hand, with dilated convolution, we can maintain constatnt stride with a wide field of vision without enhancing the computational cost of our proposed model. Finally, we can get a bigger output feature map which proved proficient for our skin lesions classification.

**Figure 4:**
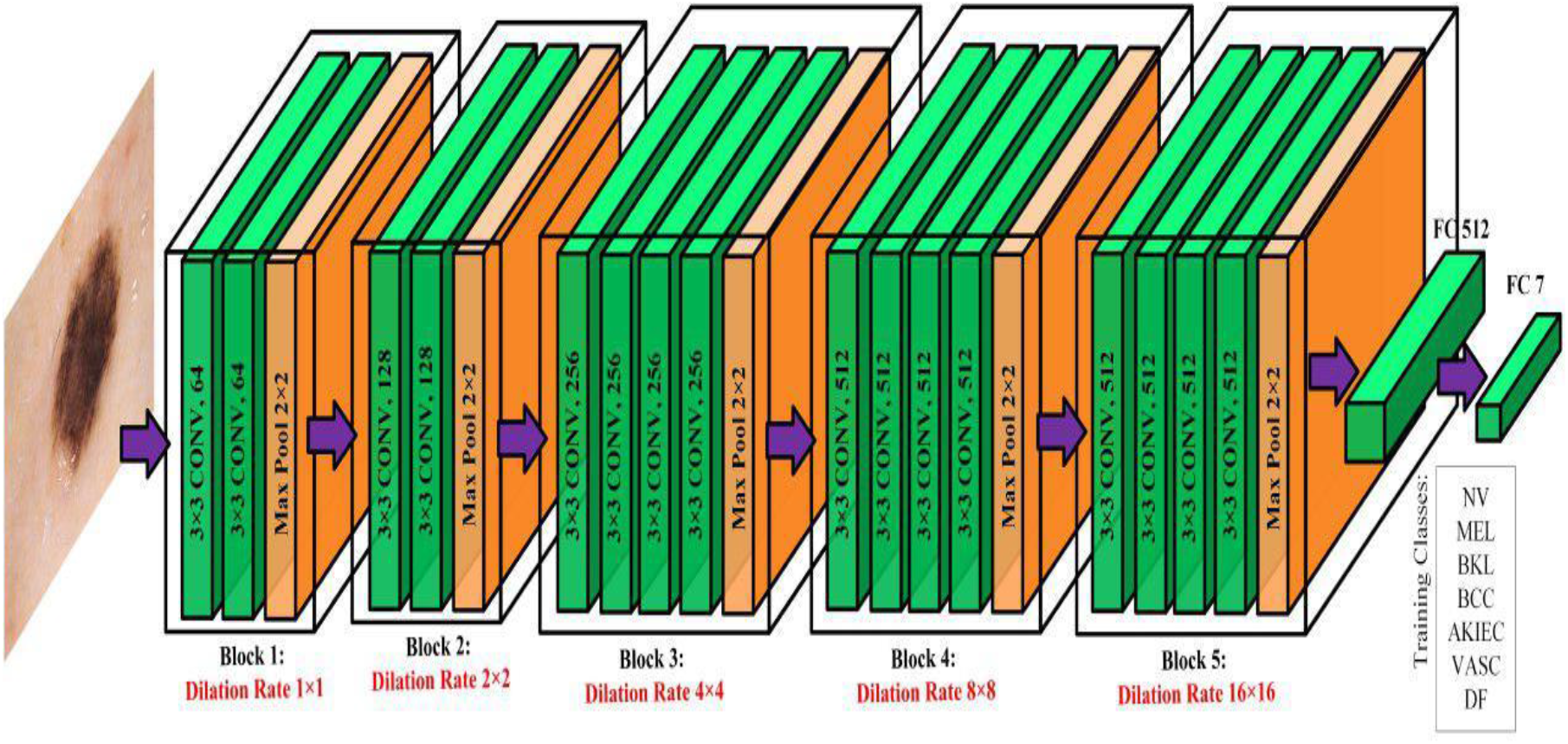
Atrous VGG16 network with different dilation rate applied on every block

The kernel or filter size of VGG16 and VGG19 is 3 × 3 in all over the model. So, the size of the receptive field will exponentially increase when we increase the value of dilation rate of our proposed methods. The most interesting fact that the number of parameters involved in this experiment is essentially alike. We can get a bigger receptive field without append any costs to models. Therefore, without improving the filter size we can get a bigger receptive field by atrous convolution which is specifically helpful when we stacked multiple dilated convolution one after another in our architectures.

Input layer shape for every proposed architecture for this research work can be referred to as *h* × *w* × *d* where, *h* = height of the image, *w* = width of the image, and *d* = for the RGB. So, the size of this layer is 192 × 256 × 3 then we initiate our work of convolutional layer and pooling layer. For both VGG networks have *dilation rate* = 1 in block one but after that, we expand the value of dilation rate as 2, 4, 8, 16 for the block two, three, four, and five respectively. We highly inspired by the multi-grid system which applies a hierarchy of grid of several different sizes [64, 65, 66, 67], some semantic image segmentation architecture [68,69] to implement this strategy. Specifically, in [69] Chen et al. choose different dilation rates within reblock4 to block7 in their proposed ResNet architecture. Seemingly, here we use VGG16 and VGG19, where block 2 to block 5 adopt distinct atrous rates respectively. Hence, we implement the standard convolutional operation for every convolutional layer throughout the model with different dilation rate.

**Figure 5:**
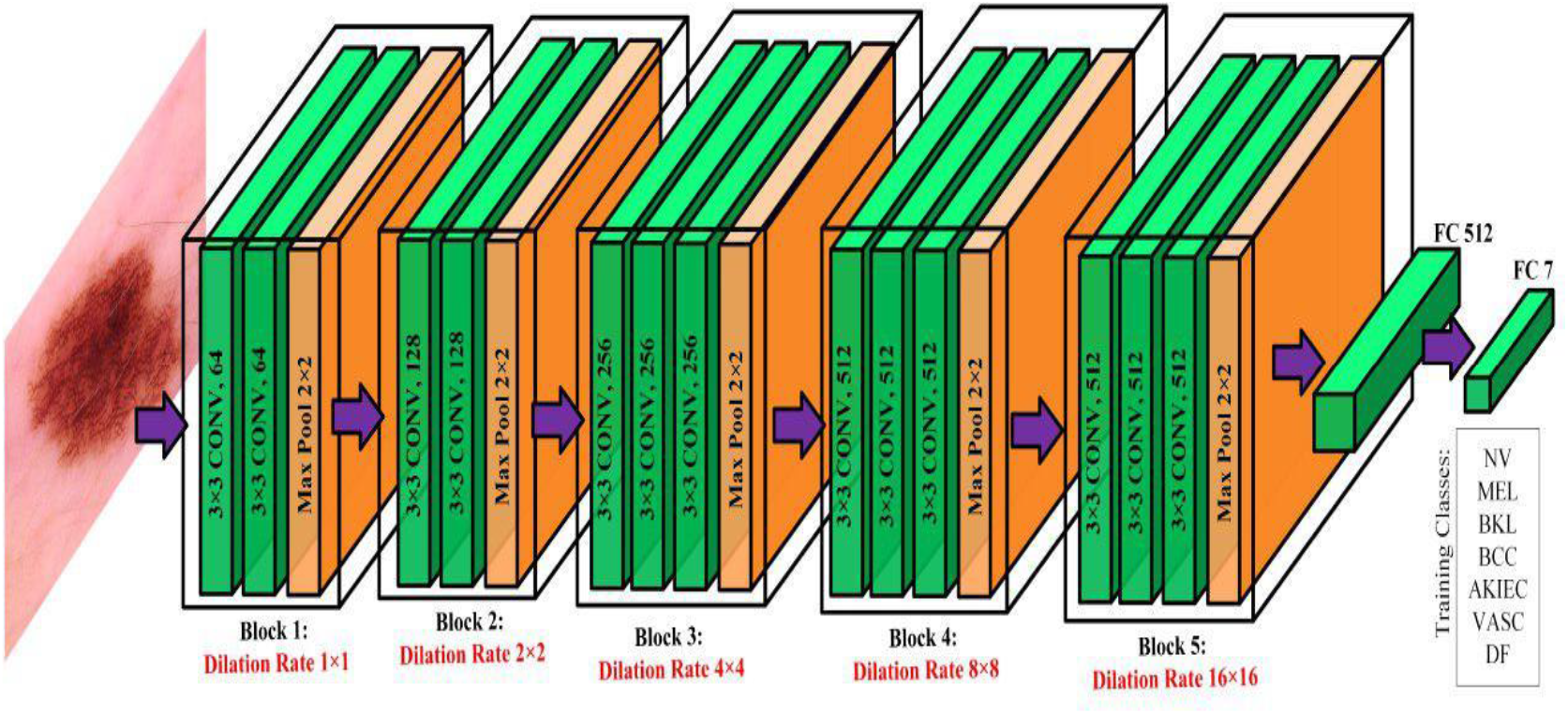
Atrous VGG19 network with different dilation rate applied on every block

Let in the standard convolutional operation 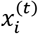 is output of layer *t* consists of 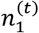 feature maps with size of 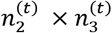. The *i*-th feature map 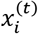 can be computed as:

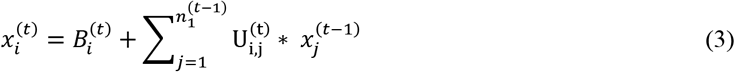

Here, bias matrix is 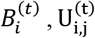 is the filter of size 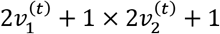 has established connection with the *j*-th feature map in the layer (*t* – 1) with *i*-th feature map in the layer.

Then we implement activation function for each convolutional layer which accomplished element-wise experiment over the input and produces output which dimension is indistinguishable to input. To be specific, activation function takes the feature map as the input and provide an activation map as the output.

So, let *t* be a non-linearity layer which take feature map 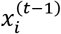 as the input from a convolutional layer (*t* – 1) and produce an output 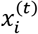 called activation volume.

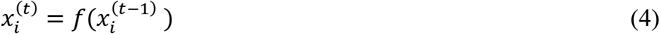

where,

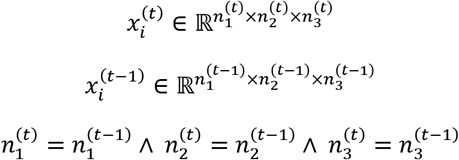

It can be denoted as, 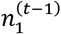 is the number of 2-d feature maps (which created by 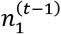 number of filters in CNN layer (*t* – 1)) each with volume 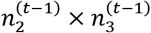.

Furthermore, we utilized rectified linear units (ReLUs) as our activation function. So, equation (4) becomes:

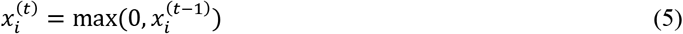

In the dilated convolution *dilation rate* > 1 provide a much larger feature map than Standard convolutional operation. In between every block of convolutional layers in VGG16 and VGG19 we have max pool technique for downsampling. Generally, it is shortening the spatial size of the activation maps created by activation function in the previous layer. It lessens the probability of overfitting and reducing computational demand progressively through the model.

Let filter *E^t^* and stride *S^t^* are two hyperparameters for pooling layer *t*. This pooling layer can take input shape 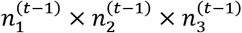 and produce a output volume 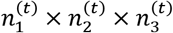. It can be denoted as:

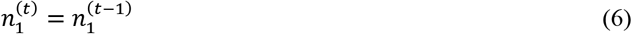

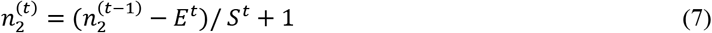

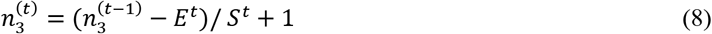

Window size *E^t^* × *E^t^* has been applied in the pooling layer.

After the fifth block of convolutional layer and fifth pooling layer, we take the feature map from fifth pooling layer and run the operation of *global max pooling* on that. It takes tensor with size *h* × *w* × *d*, here *h* × *w* known as spatial dimensions and *d* can be referred to as the number of feature maps. *Global max pooling* takes the maximal value from the spatial dimension *h* × *w* and provide an output tensor with shape 1 × 1 × *d*.

Next, we implement two fully connected layers with one dropout layer in between. The first fully connected or dense layer has 512 filters and the last one has 7 filters as we run a classification model of 7 classes. The first dense layer has “RELU” as it’s activation function and the last one has “SoftMax” activation function. If the 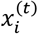 is the output vector and 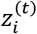 is input vector, then for layer *t* – 1 the fully connected layer can be defined:

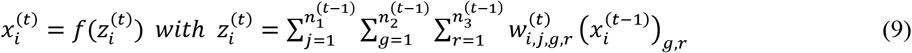

Weight parameter is 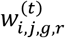 which utilize to create stochastic likelihood. In the last dense layer, we implement “SoftMax” activation function where we apply an exponential function on every element of input vector *z*. The standard “SoftMax” function 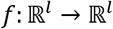 can be explained by the formula:

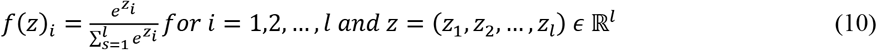

In between both dense layers, as a regularizers we use *dropout rate* = 0.50 which substantially alleviate the overfitting rate in the network but at the same time computationally inexpensive [56].

#### 3.3.2 Atrous MobileNet Architecture

MobileNet was primarily constructed to provide very low latency, small, and computationally sound architecture for embedded mobile vision application [57]. This network has three types of a convolutional layer such as a standard convolutional layer, pointwise convolutional layer, and depthwise convolutional layer. Among all the depthwise layer, five depthwise layers have a stride rate (2,2). We take these five layers to implement the atrous convolutional method. Amid these five depthwise layers, we do not change the dilation rate for the first two. The dilation rate for the first two layer remain (1,1) however, placed dilation rate (2,2) for the third and fourth depthwise layer. Furthermore, for the fifth and final depthwise convolutional layer we concatenate three depthwise convolutional 2-D layers parallelly. The dilation rate for these three layers is 4,8,16 respectively. At last, we concatenate these three layers and provide the final depthwise convolutional 2-D layer.

Basic MobileNet architecture based on depthwise separable convolutional technique where depthwise convolutional operation employs on each filter of input channel and then pointwise convolutional approach applies 1 × 1 convolution to merge the outcome of depthwise convolution. That means both filtering and combining accomplished in two steps Whereas, in the standard convolutional layer one step needed for both filtering and merging.

**Figure 6:**
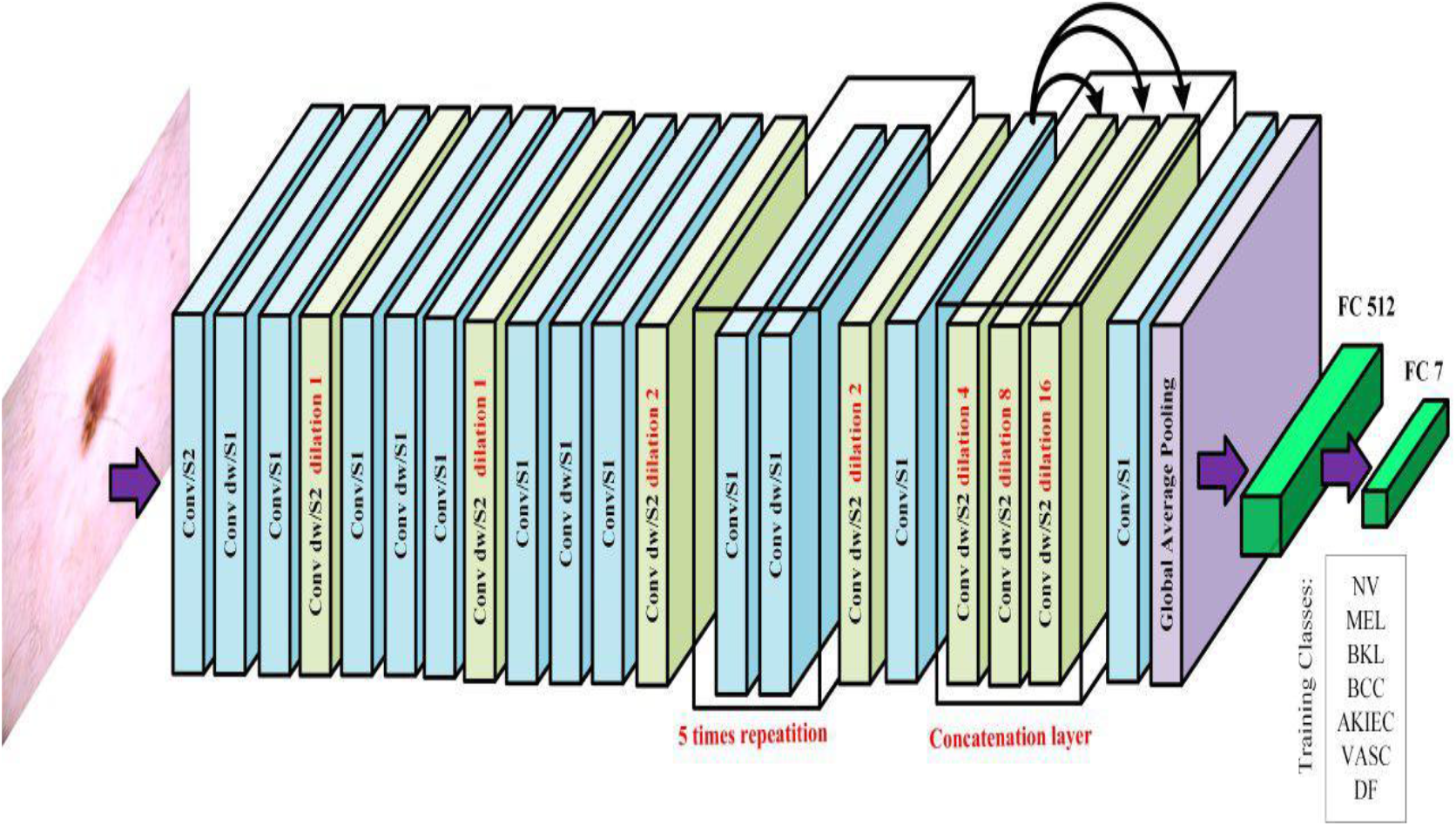
Atrous Atrous MobileNet architecture with different dilation rate on its depthwise convolutional layer which has stride (2,2). In the first two, this kind of layer has dilation rate (1,1). The 3rd and 4th layer has dilation rate (2,2). Convert three parallel depthwise layers with dilation rate (4,4), (8,8), (16,16) respectively into one depthwise layer for the final one. In the figure, dw = depthwise; S1= stride (1,1); S2 = Stride (2,2)

Let a standard convolution take input feature map *F* as *h*_1_ × *w*_1_ × *L* and provides an output feature map *G* with size *h*_2_ × *w*_2_ × *T*. Here, *h*_1_ × *w*_1_ spatial width and height for the square input feature map, *h*_2_ × *w*_2_ spatial width and height for the square output feature map, *L* is the number of input channels, and *T* is the number of output channels. So, convolutional kernel *K* can be defined as *h*_3_ × *w*_3_ × *L* × *T*, where *h*_3_ × *w*_3_ is kernel spatial dimension [57]. If the stride and padding is one, then the output feature map can be denoted as:

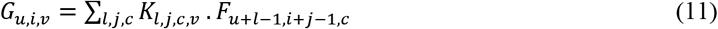

In the depthwise separable convolution this standard convolution, converted to two steps. At first, depthwise convolution applies on each filter of every input channels. Secondly, in the pointwise convolution 1 × 1 convolution employ to produce a linear combination of the product of the depthwise convolutional layer. So, the depthwise convolution can be defined as:

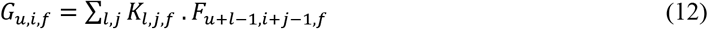

Where *K* is the kernel of depthwise convolution with the shape of *h*_3_ × *w*_3_ × *L*. Here, the *f* – *th* filter of *K* is employed to the *f* – *th* channel in F to provide filtered output feature map *G* of the *f* – *th* channel.

After that, we take the feature map from last pointwise convolutional layer and applied *global average pooling* method on the feature map. If the size of the feature map is *h* × *w* × *d* the *global average pooling* method convert the feature map into 1 × 1 × *d*, here *h* × *w* is the spatial dimension of the feature map. Additionally, *global average pooling* take the average value from the spatial dimension *h* × *w*. GAP has several advantages, for example, avoided overfitting in this layer, it shows more robust behavior to the spatial translations of the input feature map [58].

Like, VGG16 and VGG19 models in the last part we implement two fully connected layers with one dropout layer in between. The two fully connected layers have 512 filters, 7 filters respectively and the dropout rate is 0.50. Moreover, we use “SoftMax” activation function for the last fully connected layer.

#### 3.3.3 Atrous InceptionV3

From 2014, in ImageNet [59] competition the standard of neural network models achieved significant improvement because of exploiting deeper and wider architectures. Though the computational complexities still a huge issue for these wider or deeper networks. Therefore, GoogleNet [60] provided Inception network to perform competently even under the strict limitation of computation and memory budget. Additionally, Inception network also has lower computational cost than VGGNet or, other powerful networks but with high accuracy. At the same time, in the huge network, decreasing the dimensions too often is one of the main reasons for the loss of important information which is also known as “representational bottleneck”. In the InceptionV3 network, the probabilities of representational bottleneck also diminish.

There is three main focus point of InceptionV3 networks: factorizing convolution, auxiliary classifier, and coherent grid size reduction. Factorizing convolutions can be divided into three part. Firstly, factorizing into the smaller convolutions is computational sounds because in this model two 3 × 3 convolutions utilized instead of using 5 × 5 convolution to reduce the computational cost. For example, parameters for one layer of 5 × 5 filters would be (5 * 5) = 25, on the other hand two layers of 3 × 3 filters would cost (3 * 3 + 3 * 3) = 18. So, parameters decreased by 28% but it boosted the network performance. In our proposed model we call it Module A and put dilation rate (2,2) on each convolutional layer. Additionally, as we describe earlier, this strategy provides the wider receptive field with no additional computational cost and parameters. Secondly, factorize convolutions of filter shape *m* × *m* to a convolution combination of 1 × *m* and *m* × 1 convolutions. For instance, parameters number is 3*3= 9 when the filter shape is 3 × 3. Nevertheless, if we first execute a 1 × 3 convolution, then implement a 3 × 1 convolution on its outcome, the resulting number of parameters would 33% cheaper. This time the number of the parameters would be (1 * 3 + 3 * 1) = 6. We named it Inception Module B and give these layers dilation rate (2,2) like Module A to magnify the field of view with the same cost. Lastly, for high dimensional presentation, another part has been proposed in InceptionV3 architecture and we named it Inception Module C. To detach the representational bottleneck the filter banks in this module has been enlarged. In this situation, enlarged means prepare this part wider rather than deeper because if we go deeper, then we can experience an excessive dimensions reduction which can cause the loss of valuable information. We use (2,2) dilation rate for this module as well.

**Figure 7:**
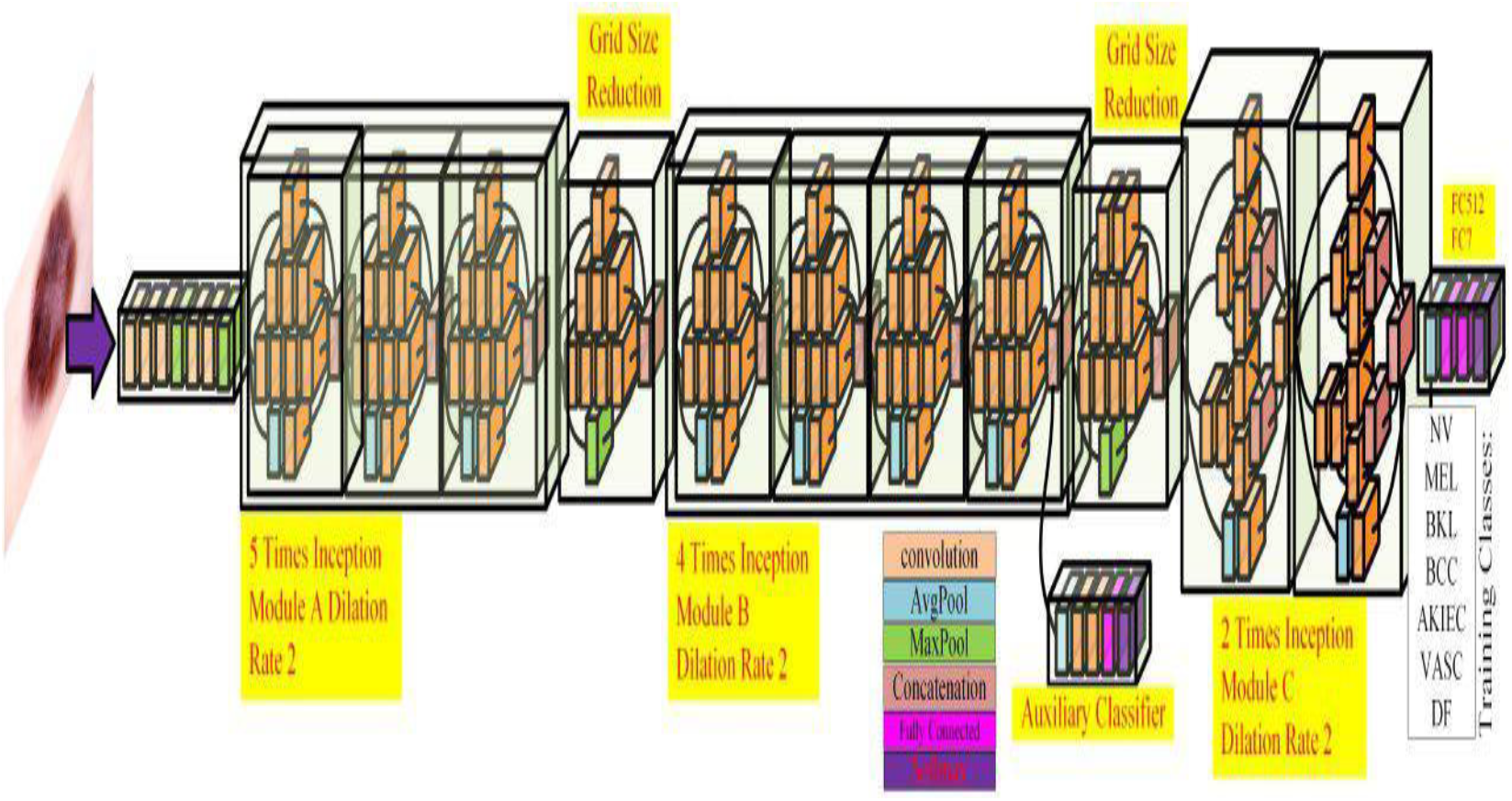
Atrous InceptionV3 network with three different modules of Inception blocks (5 times inception, 4 times Inception, 2 times inception). Every Inception block has dilation rate (2,2).

In InceptionV3, the auxiliary classifier constructed for regularization and batch normalization. Only one auxiliary classifier used in this architecture on the top of the final 17 × 17 layer. Furthermore, most of the popular deep learning architecture use max pooling to accomplish the downsizing the feature map. However, the main drawback of this procedure is max pooling can be too expensive for the architecture. Hence, in InceptionV3 in the place of max pooling, an operation called efficient grid size reduction has been applied which is less expensive and at the same time efficient for this model. In our proposed model we keep auxiliary classifier and grid size reduction sections like the basic InceptionV3.

Finally, we use *global max pooling* after the last module of this architecture. Furthermore, like previous we use two fully connected layers with filter 512, 7 respectively with dropout layer (dropout rate = 0.50) in between. Applying “RELU” as the activation function for the first dense layer and “SoftMax” for the last one.

#### 3.3.4 Transfer Learning and Fine Tune Technique

We provide the same fine tune technique for all the proposed architecture. All these architectures are pre-trained with ImageNet dataset [59] (contains approximately 1.28 million images with 1,000 class labels) and every layer of our proposed architectures has some pre-trained weight from this dataset. At first, we remove the top layers or, fully connected layers from architectures. Then add new top fully connected layers as a classifier. Top fully connected layers contain one global average pooling or, global max pooling, a fully connected layer with 512 filters, a drop out layer with 0.5 rates for regularization, and last fully connected layer with “SoftMax” activation function and seven filters. Furthermore, we freeze all layers except fully connected layer and perform feature extraction on these top fully connected layers and run until five epochs or iterations. We use this procedure to give some weight to these new fully connected layers. So, the gradient would not be too big when we start our main training part. Then we unfreeze all the layers of these proposed architectures and trained every model for 200 iterations from the very first layers to add the extra weight of HAM10000 dataset. We trained all the layers of every model in this fine-tuning technique.

## 4 Result

### 4.1 Practical Implementation

We build our main models on Keras for the frontend and used Tensorflow [44] as backend. Furthermore, use popular library Pandas for data pre-processing and Scikit Learn to see the confusion matrix, precision, recall, and F-1 score. We train every model for 200 epochs or iterations and with the mini batch size of 32. The research work performed on Intel Core i7-8750H with 4.1 GHz and an NVIDIA GeForce GTX 1050Ti GPU with 4GB GDDR5 dedicated VRAM.

We utilized Adam [70] optimizer as our optimization function with initial learning rate as 10^−4^, the exponential decay rate for the first moment estimate is 0.9, the exponential decay rate for the first moment estimates as .999, and epsilon is none. We use one callbacks functionality to reduce our learning rate during the training period when the validation loss is not reducing for 7 epochs. After 7 epochs we reduce the learning rate as the factor by (0.1)^5^. So, the new learning rate would be:

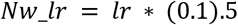

Here, *Nw_lr* = new learning rate; *lr* = present learning rate. Furthermore, the lower bound of the learning rate is 0.5e – 6.

### 4.2 Performance Analysis

#### 4.2.1 Atrous VGG16 and Atrous VGG19

Basic VGG16 and VGG19 model have overall five blocks of 2-dimensional convolutional layers and in the five blocks, each layer has dilation rate (1,1). In this experiment, we keep dilation rate (1,1) for the first block and tried numerous combinations for the other blocks (last 4 blocks) for both architecture and at last proposed which give us the best overall accuracy.

Combinations for Atrous VGG16 are as follows:

1. VGG16_Model1: {block1(dilation rate=(1,1)), block2(dilation rate=(2,2)), block3(dilation rate=(2,2)), block4(dilation rate=(2,2)), block5(dilation rate=(2,2))}
2. VGG16_Model2: {block1(dilation rate=(1,1)), block2(dilation rate=(4,4)), block3(dilation rate=(4,4)), block4(dilation rate=(4,4)), block5(dilation rate=(4,4))}
3. VGG16_Model3: {block1(dilation rate=(1,1)), block2(dilation rate=(8,8)), block3(dilation rate=(8,8)), block4(dilation rate=(8,8)), block5(dilation rate=(8,8))}
4. VGG16_Model4: {block1(dilation rate=(1,1)), block2(dilation rate=(2,2)), block3(dilation rate=(4,4)), block4(dilation rate=(8,8)), block5(dilation rate=(16,16))}

In Table 1, we show the performance analysis for basic VGG16 and several Atrous VGG16 models. Additionally, when we take dilation rate 8 for the last four layers we get accuracy lower than basic VGG16 otherwise each atrous convolutional models provide a better result than the basic VGG16. VGG16_Model4 (proposed model) acquire the highest overall accuracy for the validation set and the test set of HAM10000.

**Table 1:**
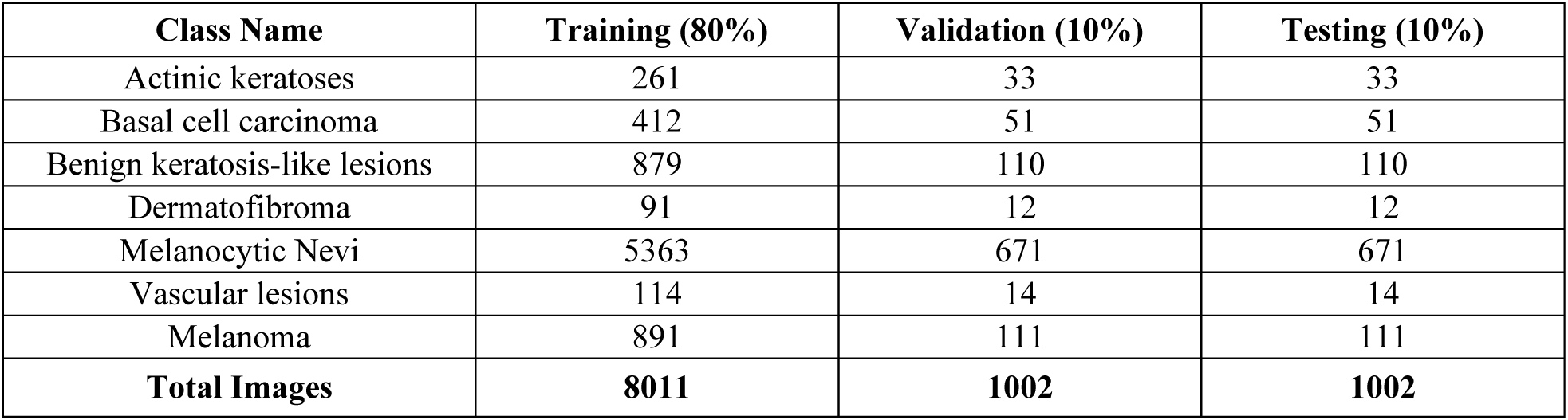
Number of images for training, validation, and test sets after splitting the HAM10000

**Table 2:** Comparison of several combinations of Atrous VGG16 and Basic VGG16 for overall accuracy

| Model Name | Peak Validation Accuracy (%) | Test Accuracy (%) |
| --- | --- | --- |
| Basic VGG16 | 85.21 | 84.03 |
| VGG16_Model1 | 86.38 | 85.47 |
| VGG16_Model2 | 87.09 | 86.40 |
| VGG16_Model3 | 83.94 | 83.32 |
| VGG16_Model4 (Proposed Method) | **90.10** | **87.42** |

Combinations for Atrous VGG19 are as follows:

1. VGG19_Model1: {block1(dilation rate=(1,1)), block2(dilation rate=(2,2)), block3(dilation rate=(2,2)), block4(dilation rate=(2,2)), block5(dilation rate=(2,2))}
2. VGG19_Model2: {block1(dilation rate=(1,1)), block2(dilation rate=(4,4)), block3(dilation rate=(4,4)), block4(dilation rate=(4,4)), block5(dilation rate=(4,4))}
3. VGG19_Model3: {block1(dilation rate=(1,1)), block2(dilation rate=(8,8)), block3(dilation rate=(8,8)), block4(dilation rate=(8,8)), block5(dilation rate=(8,8))} VGG19_Model4: {block1(dilation rate=(1,1)), block2(dilation rate=(2,2)), block3(dilation rate=(4,4)), block4(dilation rate=(8,8)), block5(dilation rate=(16,16))}

In Table 3, we display the comparison of basic VGG19 and several Atrous VGG19 models. In addition, except one every atrous convolution methods exhibit a better result than the basic model. Overall, VGG19_Model4 (proposed model) obtain the top accuracy for both the validation set and the test set of HAM10000.

**Table 3:** Comparison of several combinations of Atrous VGG19 and Basic VGG19

| Model Name | Peak Validation Accuracy | Test Accuracy |
| --- | --- | --- |
| Basic VGG19 | 84.02 | 82.96 |
| VGG19_Model1 | 85.82 | 84.84 |
| VGG19_Model2 | 85.79 | 84.60 |
| VGG19_Model3 | 82.95 | 82.37 |
| VGG19_Model4 (Proposed Method) | **86.39** | **85.02** |

#### 4.2.2 Atrous MobileNet

There is a total of five depth-wise 2D convolutional layers which has stride (2,2) in the MobileNet network and the dilation rate for each of these layers is (1,1). In this research work, we retain the dilation rate (1,1) for the first two depth-wise convolutional layers and tested different combinations for the last three layers.

Combinations for Atrous MobileNet are as follows:

1. MobileNet_Model1: {depthwise_Conv_Layer1(dilation rate(1,1)), depthwise_Conv_Layer2(dilation rate(1,1)), depthwise_Conv_Layer3(dilation rate(2,2)), depthwise_Conv_Layer4(dilation rate(2,2)), depthwise_Conv_Layer5(dilation rate(2,2))}
2. MobileNet_Model2: {depthwise_Conv_Layer1(dilation rate(1,1)), depthwise_Conv_Layer2(dilation rate(1,1)), depthwise_Conv_Layer3(dilation rate(4,4)), depthwise_Conv_Layer4(dilation rate(4,4)), depthwise_Conv_Layer5(dilation rate(4,4))}
3. MobileNet_Model3: {depthwise_Conv_Layer1(dilation rate(1,1)), depthwise_Conv_Layer2(dilation rate(1,1)), depthwise_Conv_Layer3(dilation rate(8,8)), depthwise_Conv_Layer4(dilation rate(8,8)), depthwise_Conv_Layer5(dilation rate(8,8))}
4. MobileNet_Model4: [depthwise_Conv_Layer1(dilation rate(1,1)), depthwise_Conv_Layer2(dilation rate=(1,1)), depthwise_Conv_Layer3(dilation rate(2,2)), depthwise_Conv_Layer4(dilation rate(2,2)), depthwise_Conv_Layer5 {Concatenate(depthwise_Conv(dilation rate(4,4), depthwise_Conv(dilation rate(8,8))}]
5. MobileNet_Model5: [depthwise_Conv_Layer1(dilation rate(1,1)), depthwise_Conv_Layer2(dilation rate=(1,1)), depthwise_Conv_Layer3(dilation rate(2,2)), depthwise_Conv_Layer4(dilation rate(2,2)), depthwise_Conv_Layer5 {Concatenate(depthwise_Conv(dilation rate(4,4), depthwise_Conv(dilation rate(8,8), depthwise_Conv(dilation rate(16,16))}]

In Table 4, we present the contrast of basic MobileNet and several Atrous MobileNet methods. Interestingly, when we take the dilation rate (4,4) and (8,8) for the last three layers then the model delivers less accuracy than the basic MobileNet. Overall, MobileNet_Model5 (proposed model) procure the best accuracy for both the validation set and the test set of HAM10000.

**Table 4:** Comparison of several combinations of Atrous MobileNet and Basic MobileNet for overall accuracy

| Model Name | Peak Validation Accuracy (%) | Test Accuracy (%) |
| --- | --- | --- |
| Basic MobileNet | 87.09 | 85.87 |
| MobileNet_Model1 | 87.90 | 86.76 |
| MobileNet_Model2 | 86.84 | 85.62 |
| MobileNet_Model3 | 85.57 | 84.42 |
| MobileNet_Model4 | 88.73 | 87.31 |
| MobileNet_Model5 (Proposed Method) | **89.48** | **88.22** |

#### 4.2.3 Atrous InceptionV3

There are three sets of inception module in InceptionV3 network, for instance, 5×Inception Module A, 4×Inception Module B, and 2×Inception Module C. These Module’s 2-dimensional convolutional layers have dilation rate (1,1). So, we change the dilation rate of these convolutional layers and inspect the performance analysis for various combinations. There are two Grid Size Reduction blocks and one Auxiliary Classifier block. In addition, we leave them as like they were before.

Combinations for Atrous InceptionV3 are as follows:

1. InceptionV3_Model1: {Module A(dilation rate(2,2)), Module B (dilation rate(2,2)), Module C (dilation rate(2,2))}
2. InceptionV3_Model2: {Module A(dilation rate(4,4)), Module B (dilation rate(4,4)), Module C (dilation rate(4,4))}
3. InceptionV3_Model3: {Module A(dilation rate(8,8)), Module B (dilation rate(8,8)), Module C (dilation rate(8,8))}
4. InceptionV3_Model4: {Module A(dilation rate(2,2)), Module B (dilation rate(4,4)), Module C (dilation rate(8,8))}

In Table 5, we exhibit the accuracy comparison of numerous Atrous InceptionV3 network and basic InceptionV3 model. When the dilation rate (2,2) for three Modules the InceptionV3 produce the best result for the validation set and the test set among all the models. However, the performance decline when we increase the dilation rate to (4,4) and (8,8). Dilation rate (8,8) for the three modules even produce low accuracy than the basic model.

**Table 5:** Comparison for several combination of Atrous InceptionV3 and Basic InceptionV3

| Model Name | Peak Validation Accuracy | Test Accuracy |
| --- | --- | --- |
| Basic InceptionV3 | 87.62 | 86.72 |
| InceptionV3_Model1 (Proposed Model) | **90.95** | **89.81** |
| InceptionV3_Model2 | 89.56 | 88.12 |
| InceptionV3_Model3 | 86.18 | 85.30 |
| InceptionV3_Model4 | 87.71 | 87.02 |

### 4.3 Evaluation and Comparison of Four Proposed Models

The HAM10000 is hugely class imbalanced dataset. Therefore, we wanted to see the performance of for recall, precision, and F1 score for each individual proposed architecture. In Table 5, Table 6, Table 7, Table 8 we show the recall, precision, f1-score, micro average, macro average, and weighted average of VGG16_Model4, VGG19_Model4, MobileNet_Model4, InceptionV3_Model4 respectively for the test set to see how each class performed on our proposed architectures.

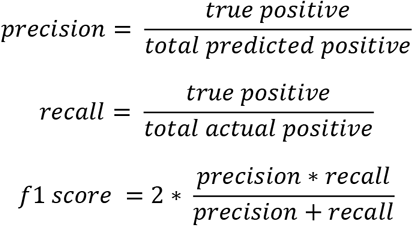

**Table 6:** Precision, recall, F1-score, micro avg., macro avg., and weighted avg. for the test set of HAM10000 dataset of proposed VGG16 model (VGG16_Model4)

| Class Name | Precision | Recall | F1-Score |
| --- | --- | --- | --- |
| Actinic keratoses | 0.64 | 0.42 | 0.51 |
| Basal cell carcinoma | 0.80 | 0.84 | 0.82 |
| Benign keratosis like lesions | 0.72 | 0.76 | 0.74 |
| Dermatofibroma | 0.67 | 0.67 | 0.67 |
| Melanocytic Nevi | 0.94 | 0.96 | 0.95 |
| Vascular lesions | 1.00 | 0.79 | 0.88 |
| Melanoma | 0.72 | 0.66 | 0.69 |
| **Micro Avg** | 0.87 | 0.87 | 0.87 |
| **Macro Avg** | 0.78 | 0.73 | 0.75 |
| **Weighted Avg** | 0.87 | 0.87 | 0.87 |

**Table 7:** Precision, recall, F1-score, micro avg., macro avg., and weighted avg. for the test set of HAM10000 dataset of proposed VGG19 model (VGG19_Model4)

| Class Name | Precision | Recall | F1-Score |
| --- | --- | --- | --- |
| Actinic keratoses | 0.61 | 0.58 | 0.59 |
| Basal cell carcinoma | 0.79 | 0.65 | 0.71 |
| Benign keratosis like lesions | 0.66 | 0.82 | 0.73 |
| Dermatofibroma | 0.67 | 0.33 | 0.44 |
| Melanocytic Nevi | 0.93 | 0.93 | 0.93 |
| Vascular lesions | 0.91 | 0.71 | 0.80 |
| Melanoma | 0.67 | 0.65 | 0.66 |
| **Micro Avg** | 0.85 | 0.85 | 0.85 |
| **Macro Avg** | 0.75 | 0.67 | 0.70 |
| **Weighted Avg** | 0.85 | 0.85 | 0.85 |

**Table 8:** Precision, recall, F1-score, micro avg., macro avg., and weighted avg. for the test set of HAM10000 dataset of proposed MobileNet architecture (MobileNet_Model5)

| Class Name | Precision | Recall | F1-Score |
| --- | --- | --- | --- |
| Actinic keratoses | 0.79 | 0.45 | 0.58 |
| Basal cell carcinoma | 0.94 | 0.65 | 0.77 |
| Benign keratosis like lesions | 0.65 | 0.86 | 0.74 |
| Dermatofibroma | 0.90 | 0.75 | 0.82 |
| Melanocytic Nevi | 0.96 | 0.94 | 0.95 |
| Vascular lesions | 0.92 | 0.79 | 0.85 |
| Melanoma | 0.73 | 0.79 | 0.76 |
| **Micro Avg** | 0.88 | 0.88 | 0.88 |
| **Macro Avg** | 0.84 | 0.75 | 0.78 |
| **Weighted Avg** | 0.89 | 0.88 | 0.88 |

**Table 9:**
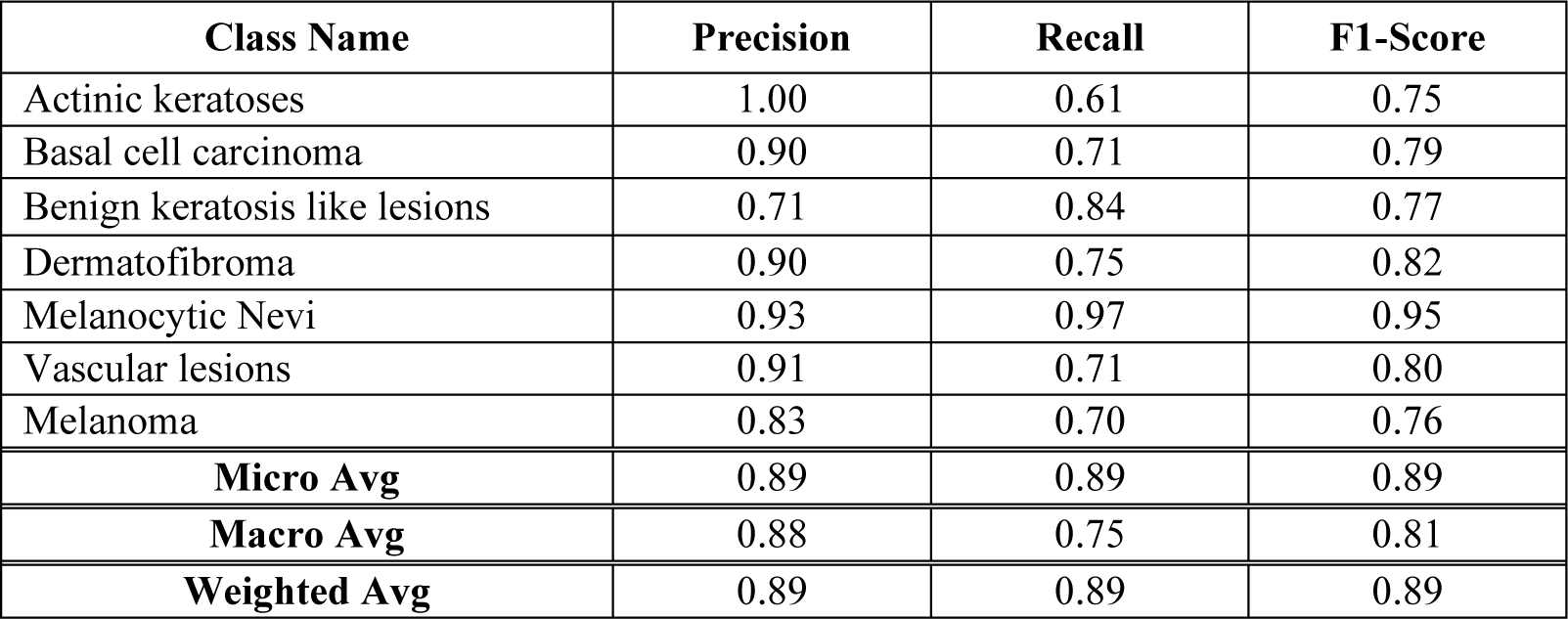
Precision, recall, F1-score, micro avg., macro avg., and weighted avg. for the test set of HAM10000 dataset of proposed InceptionV3 architecture (InceptionV3_Model1)

Precision, or positive predictive value means the percentage of our outcomes which are pertinent (how many selected items are relevant?). Recall, sensitivity, or true positive rate means the percentage of total relevant outcomes accurately classified the selected model (how many relevant items are selected?). F1-score is the harmonic mean of the precision and recall (measure of the test accuracy). For example, if we consider the case of Melanoma, then true positive means the number of accurately predicted Melanoma lesion by the model. Precision refers to how many of them are actual positive out of those total predicted positive. In this example, the total actual positive means the number of Melanoma lesion images exist in the test set.

### 4.4 Confusion Matrix of Four Proposed Atrous Convolutional Models

In table 10, 11, 12 and 13 we demonstrate the confusion matrix value of HAM10000 dataset for our four proposed atrous convolutional neural network architectures. Confusion matrix used to summarize the performance of the architecture. Only classification accuracy is not enough to see the performance of our proposed algorithm on HAM10000 because it is a class imbalanced dataset. Confusion matrix can give a clear idea of what the classification architecture getting right and what kinds of mistakes it made. For example, in the first row of table 10 VGG16_Model4 correctly labeled 14 images as Actinic Keratoses, however, it mistakenly labeled 5 Actinic Keratoses images as Basal Cell Carcinoma, 8 images as Benign Keratosis like lesions, 1 image as Dermatofibroma, and 5 images as Melanoma. Similarly, in row 5 the proposed VGG16 model accurately classified 643 images as Melanocytic Nevi but it wrongly assumes 1 image as Actinic keratoses, 1 image as Basal cell carcinoma, 15 images as Benign keratosis like lesions, and 11 images as Melanoma. Melanocytic Nevi is the biggest class in this test set. On the other hand, Dermatofibroma has a minimal number of images. The confusion matrix size is 7×7 because we have seven classes in our dataset. For these seven classes proposed VGG16, MobileNet, and InceptionV3 displayed better result than proposed VGG19. Among these four dilated models, dilated IncaptionV3 provide the superior result for Actinic keratoses, and Melanocytic Nevi. For Melanoma, and Benign keratosis like lesions, MobileNet shows the impressive outcome. Furthermore, atrous VGG16 is the triumph for Basal cell carcinoma. Proposed VGG16 and MobileNet yield higher accuracy for Vascular lesions. Lastly, for Dermatofibroma both InceptionV3 and MobileNet present better result than both proposed VGG networks.

**Table 10:** Confusion Matrix for proposed VGG16 model (VGG16_Model4)

|  |  | Predicted Class |  |  |  |  |  |  |
| --- | --- | --- | --- | --- | --- | --- | --- | --- |
|  |  | Actinic<br>keratoses | Basal cell<br>carcinom<br>a | Benign<br>keratos<br>is like<br>lesions | Dermat<br>ofibro<br>ma | Melanoc<br>ytic<br>Nevi | Vascular<br>lesions | Melanoma |
| Actual<br>Class or,<br>True<br>Class | Actinic<br>keratoses | **14** | 5 | 8 | 1 | 0 | 0 | 5 |
|  | Basal cell<br>carcinoma | 2 | **43** | 1 | 1 | 2 | 0 | 2 |
|  | Benign<br>keratosis like<br>lesions | 2 | 1 | **84** | 1 | 13 | 0 | 9 |
|  | Dermatofibro<br>ma | 0 | 1 | 1 | **8** | 1 | 0 | 1 |
|  | Melanocytic<br>Nevi | 1 | 1 | 15 | 0 | **643** | 0 | 11 |
|  | Vascular<br>lesions | 0 | 1 | 0 | 0 | 2 | **11** | 0 |
|  | Melanoma | 3 | 2 | 7 | 1 | 25 | 0 | **73** |

**Table 11:**
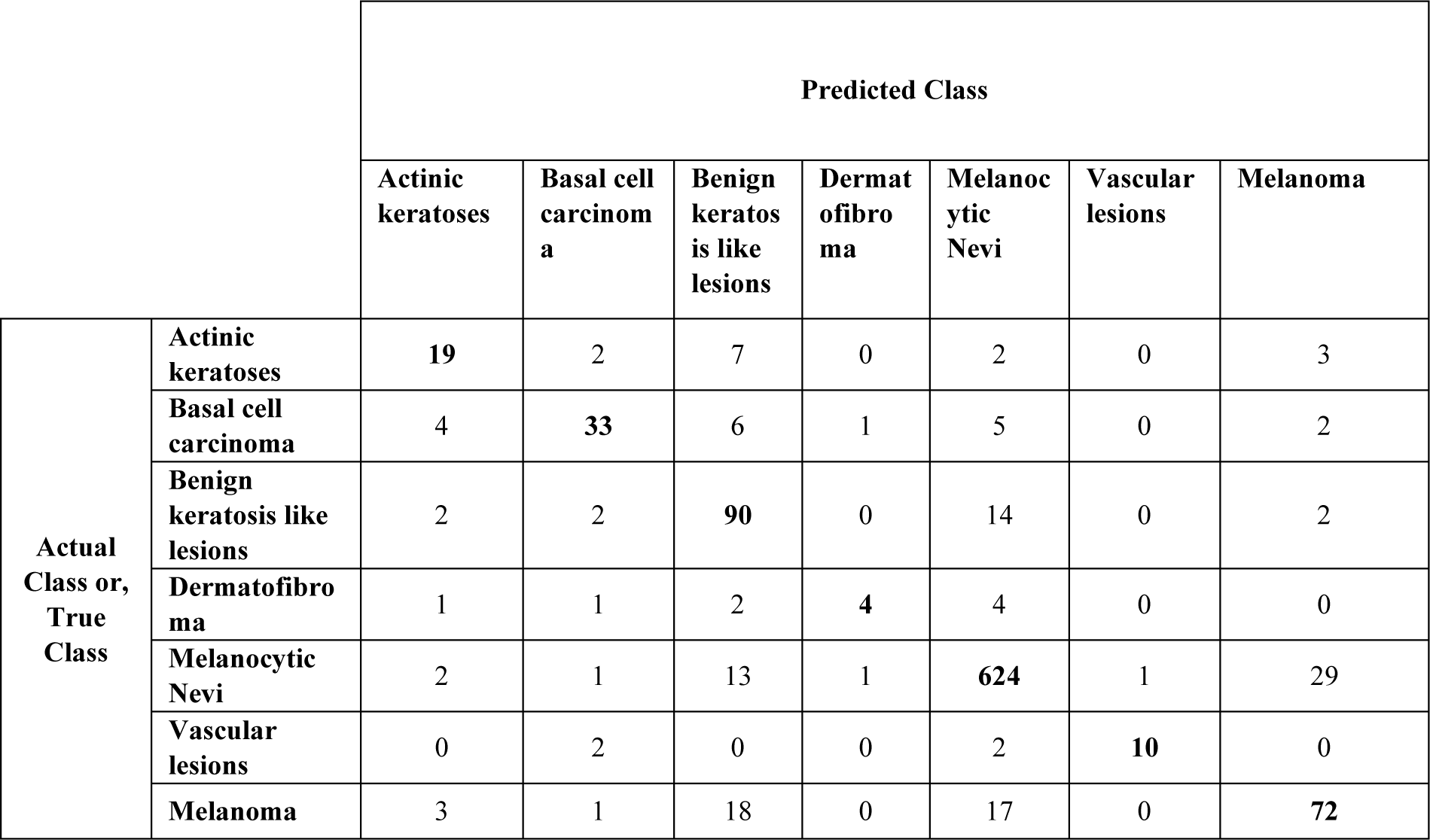
Confusion Matrix for proposed VGG19 model (VGG19_Model4)

**Table 12:** Confusion Matrix for proposed MobileNet architecture (MobileNet_Model5)

|  |  | Predicted Class |  |  |  |  |  |  |
| --- | --- | --- | --- | --- | --- | --- | --- | --- |
|  |  | Actinic<br>keratoses | Basal cell<br>carcinom<br>a | Benign<br>keratos<br>is like<br>lesions | Dermat<br>ofibro<br>ma | Melanoc<br>ytic<br>Nevi | Vascular<br>lesions | Melanoma |
| Actual<br>Class or,<br>True<br>Class | Actinic<br>keratoses | 15 | 0 | 15 | 0 | 0 | 0 | 3 |
|  | Basal cell<br>carcinoma | 2 | 33 | 11 | 1 | 3 | 0 | 1 |
|  | Benign<br>keratosis like<br>lesions | 1 | 1 | 95 | 0 | 6 | 1 | 6 |
|  | Dermatofibro<br>ma | 1 | 0 | 1 | 9 | 1 | 0 | 0 |
|  | Melanocytic<br>Nevi | 0 | 0 | 16 | 0 | 633 | 0 | 22 |
|  | Vascular<br>lesions | 0 | 1 | 1 | 0 | 1 | 11 | 0 |
|  | Melanoma | 0 | 0 | 13 | 0 | 10 | 0 | 88 |

**Table 13:** Confusion Matrix for proposed InceptionV3 architecture (InceptionV3_Model1)

|  |  | Predicted Class |  |  |  |  |  |  |
| --- | --- | --- | --- | --- | --- | --- | --- | --- |
|  |  | Actinic<br>keratoses | Basal cell<br>carcinom<br>a | Benign<br>keratos<br>is like<br>lesions | Dermat<br>ofibro<br>ma | Melanoc<br>ytic<br>Nevi | Vascular<br>lesions | Melanoma |
| Actual<br>Class or,<br>True<br>Class | Actinic<br>keratoses | 20 | 1 | 6 | 0 | 2 | 0 | 4 |
|  | Basal cell<br>carcinoma | 0 | 36 | 7 | 1 | 6 | 0 | 1 |
|  | Benign<br>keratosis like<br>lesions | 0 | 1 | 92 | 0 | 15 | 0 | 2 |
|  | Dermatofibro<br>ma | 0 | 0 | 1 | 9 | 1 | 0 | 1 |
|  | Melanocytic<br>Nevi | 0 | 1 | 13 | 0 | 648 | 1 | 8 |
|  | Vascular<br>lesions | 0 | 1 | 0 | 0 | 3 | 10 | 0 |
|  | Melanoma | 0 | 0 | 10 | 0 | 23 | 0 | 78 |

**Table 14:** Model complexity comparison of four proposed Atrous Convolutional architecture

| Model Name | Parameters |
| --- | --- |
| Proposed VGG16 | 14,980,935 (14.98 million) |
| Proposed VGG19 | 20,290,631 (20.29 million) |
| Proposed MobileNet | **3,757,255** (3.76 million) |
| Proposed InceptionV3 | 22,855,463 (22.85 million) |

**Table 15:** Performance analysis with different input image size

| Input Image Size ($W \times H$) | Atrous VGG16 (%) | Atrous VGG19 (%) | Proposed InceptionV3 (%) |
| --- | --- | --- | --- |
| $64 \times 64$ | 79.52 | 76.41 | 84.98 |
| $128 \times 96$ | 85.93 | 82.71 | 87.57 |
| $256 \times 192$ | **87.42** | 84.92 | **89.81** |
| $320 \times 320$ | 86.45 | **85.02** | 88.16 |

**Table 16:** performance analysis with different input image shape for Atrous MobileNet

| Input Image Size ($W \times H$) | Atrous MobileNet (%) |
| --- | --- |
| $128 \times 128$ | 83.37 |
| $160 \times 160$ | 86.60 |
| $192 \times 192$ | 87.85 |
| $224 \times 224$ | **88.22** |

### 4.5 Model Complexity

In this section, we exhibit the model complexity for our four proposed architecture. Though proposed InceptionV3 network provides overall best accuracy among these four models, it uses more parameters than any other proposed model. Thus, this model’s computational complexity is superior too. On the other hand, MobileNet produces less computational complexity (only 3.75 million).

### 4.6 Performance Analysis Based on Image Size

We examine several combinations of image sizes to inspect the overall outcome (test accuracy) of our proposed models. These image size combinations for Atrous VGG16, VGG19, and InceptionV3 are 64×64,128×96, 256×192, 320×320.

The accuracy vigorously improved when we enhance the image dimension from 64×64 to 128×96. Furthermore, it continuously improving the test result up to the image size 256×192 and for this combination, it provides the best outcome for every model except VGG19. This performance depletion happened because when we enhance the image size after a certain point the models cannot able to handle the training challenge of these quadruple number of image pixels [59]. Vast architecture with more layers and capacity can perform better for the high-resolution images. Furthermore, high-resolution image takes more time to train the whole model and that’s why when we take 320×320 image size it increases the time complexity of the models. Therefore, we think that 256×192 image size yielded high accuracy with favorable training time for every model.

Image shape combinations for Atrous MobileNet are: (128×128), (160×160), (192×192), (224×224). We utilized four different combinations to check the accuracy of MobileNet but get highest accuracy with 224×224 image resolution.

## 5 Conclusion and Future Work

Skin cancer is one of the most frequent and threatening kinds of cancer not only in North America but also all over the world. The recent advancement of deep convolutional neural network in medical imaging gives us the inspiration to classify skin lesions from dermoscopic skin images. We implement atrous or, dilated convolution in four of most popular deep neural network architectures to establish an automated model to classify several classes of skin lesion. We started with a proof of concept about the classification of skin lesions. To make the dataset, HAM10000, more suitable for our CNN models, we applied many data preprocessing and data augmentation methods. We have tested four proposed convolutional neural networks (CNN) which are atrous VGG16, atrous VGG19, atrous MobileNet, and atrous InceptionV3. All of these methods provide higher accuracy than the basic architectures and improved recall, precision, and F-1 score. We achieved 87.42%, 85.02%, 88.22%, and 89.81% overall accuracy for our proposed VGG16, VGG19, MobileNet, and InceptionV3 respectively. Our models provide better results than any state-of-the-art methods on skin cancer classification in terms of number of classes considering image noise presence and class imbalance. Among them, dilated InceptionV3 shows best classification accuracy while dilated MobileNet has the lightest computational complexities.

We have still several limitations to tackle. For example, without a powerful GPU setup, each model could take several days to accomplish the whole training and testing process. So, the time complexity still a big issue in this training procedure. Besides, we want to observe the result for some other vast neural network model such as DenseNet. We still fancy to achieve more overall accuracy and per-class accuracy for our presented methods. Next, every of our proposed models has been pre-trained with ImageNet dataset but ImageNet dataset is not well established for skin images. We use the fine tuning technique to give every layer the extra weight of HAM10000. So, we have to train every layer of each model to give them this extra weight which enhances the space complexities and time complexities. Therefore, we like to minimize these complexities for our proposed models and want to prepare these models for practical medical applications.

